# Phylogenetic and functional analyses of *N*^6^-methyladenosine RNA methylation factors in the wheat scab fungus *Fusarium graminearum*

**DOI:** 10.1101/2023.08.11.552984

**Authors:** Hyeonjae Kim, Jianzhong Hu, Hunseung Kang, Wonyong Kim

**Author notes:** Address correspondence to Wonyong Kim.

## Abstract

In eukaryotes, *N*^6^-methyladenosine (m^6^A) RNA modification plays crucial roles in governing the fate of RNA molecules and has been linked to various developmental processes. However, the phyletic distribution and functions of genetic factors responsible for m^6^A modification remain largely unexplored in fungi. To get insights into evolution of m^6^A machineries, we reconstructed global phylogenies of potential m^6^A writers, readers, and erasers in fungi. Substantial copy number variations were observed, ranging from up to five m^6^A writers in early-diverging fungi to a single copy in the subphylum Pezizomycotina, which primarily comprises filamentous fungi. To characterize m^6^A factors in a phytopathogenic fungus *Fusarium graminearum*, we generated knockout mutants lacking potential m^6^A factors including the sole m^6^A writer *MTA1*. However, the resulting knockouts did not exhibit any noticeable phenotypic changes during vegetative and sexual growth stages. As obtaining a homozygous knockout lacking *MTA1* was likely hindered by its essential role, we generated *MTA1*-overexpressing strains (*MTA1*-OE). The *MTA1*-OE5 strain showed delayed conidial germination and reduced hyphal branching, suggesting its involvement during vegetative growth. Consistent with these findings, the expression levels of *MTA1* and a potential m^6^A reader *YTH1* were dramatically induced in germinating conidia, followed by the expression of potential m^6^A erasers at later vegetative stages. Several genes including transcription factors, transporters and various enzymes were found to be significantly up- and down-regulated in the *MTA1*-OE5 strain. Overall, our study highlights the functional importance of the m^6^A methylation during conidial germination in *F. graminearum* and provides a foundation for future investigations into m^6^A modification sites in filamentous fungi.

**Importance:** *N*^6^-methyladenosine (m^6^A) RNA methylation is a reversible posttranscriptional modification that regulates RNA function and plays a crucial role in diverse developmental processes. This study addresses the knowledge gap regarding phyletic distribution and functions of m^6^A factors in fungi. The identification of copy number variations among fungal groups enriches our knowledge regarding the evolution of m^6^A machinery in fungi. Functional characterization of m^6^A factors in a phytopathogenic filamentous fungus *Fusarium graminearum* provides insights into the essential role of the m^6^A writer *MTA1* in conidial germination and hyphal branching. The observed effects of overexpressing *MTA1* on fungal growth and gene expression patterns of m^6^A factors throughout the life cycle of *F. graminearum* further underscore the importance of m^6^A modification in conidial germination. Overall, this study significantly advances our understanding of m^6^A modification in fungi, paving the way for future research into its roles in filamentous growth and potential applications in disease control.

## Introduction

Eukaryotic RNA undergoes over 100 chemical modifications that can impact RNA processing and metabolism (1–3). These modifications include mRNA capping, mRNA polyadenylation, RNA splicing, and RNA methylation (4). Among various RNA methylation, *N*^6^-methyladenosine (m^6^A) is characterized by the methylation of the sixth nitrogen atom on adenosine within RNA. m^6^A was first described in mammalian cells fifty years ago, and is the most prevalent and abundant modification found in eukaryotic mRNA (5, 6). This modification is reversible and regulated by a set of enzymes that function as writers (methyltransferases), erasers (demethylases), and readers (RNA-binding proteins that recognize m^6^A) (7). Due to its reversible nature, m^6^A modification serves as a rapid response to environmental stress and regulates various processes including development, immune reactions, and cancer progression in animals, fungi and plants (8–15).

m^6^A RNA modification is estimated to occur in approximately 0.1–0.4% of adenosine nucleotides found in mammalian mRNAs (16, 17). In mammals, the methyltransferase complex responsible for m^6^A methylation includes proteins, such as methyltransferase-like protein 3 (METTL3), <u>met</u>hyltransferase-like protein <u>14</u> (METTL 14), and the pre-mRNA-splicing regulator <u>W</u>ilms tumor 1-<u>a</u>ssociated <u>p</u>rotein (WTAP) (18–20). METTL3 serves as the catalytic subunit, forming a heterodimer with its paralogue, METTL14. WTAP is a regulatory subunit whose function is to recruit the m^6^A methyltransferase complex to the target mRNA in nuclear speckles, and is believed to act as a bridge between the METTL3/ METTL14 heterodimer and accessory proteins (21). Among the accessory proteins, <u>vir</u>ilizer-like <u>m</u>ethyltransferase-<u>a</u>ssociated protein (VIRMA) is implicated in stabilizing the methyltransferase complex, and plays a role in the selection of specific sites for the m^6^A modification (22). <u>YT</u>521-B <u>h</u>omology (YTH)-domain proteins are known as m^6^A readers located in the cytoplasm that influence on translation of methylated mRNAs and their subsequent degradation (23, 24). In humans, the <u>YTH d</u>omain family <u>2</u> protein (YTHDF2) is involved in controlling mRNA stability by specifically binding to m^6^A-modified mRNAs and relocating them from ribosomes to processing bodies (25). Demethylation of m^6^A is catalyzed by 2-oxoglutarate and iron-dependent [2OG-Fe(II)] dioxygenases AlkB-like domain-containing proteins and the fat mass and obesity associated protein in mammals (26, 27). These diverse m^6^A factors collectively contribute to m^6^A RNA modification, playing a significant role in RNA metabolism and gene regulation.

In the budding yeast *Saccharomyces cerevisiae*, the m^6^A writer IME4 (Inducer of <u>me</u>iosis <u>4</u>) is homologous to human METTL3 that catalyzes m^6^A modification of specific target RNAs involved in the initiation of sexual development and sporulation (28). *IME4* is required for proper entry into sexual development and progression through meiotic divisions in diploid cells, and is also known to regulate triacylglycerol metabolism, vacuolar morphology and mitochondrial morphology in haploid cells (29, 30). An yeast two-hybrid screening identified core components of the methyltransferase complex composed of two catalytic factors IME4 and KAR4 (<u>Kar</u>yogamy <u>4</u>, orthologous to human METTL14), as well as two non-catalytic factors MUM2 (<u>Mu</u>ddled <u>m</u>eiosis <u>2</u>, orthologous to human WTAP), and SLZ1 (<u>S</u>porulation leucine <u>z</u>ipper <u>1</u> orthologous to human ZCH13H3) in the budding yeast (31). Recent work revealed that an interacting protein with KAR4, Ygl036wp (thereafter, coined VIR1), in the budding yeast shares a folding pattern similar to VIRMA, despite lack of discernable protein domains (32, 33). In the budding yeast, SLZ1 enables the methyltransferase complex comprising IME4, KAR4, MUM2 (WTAP), and VIR1 to function in m^6^A deposition (33).

The genome of the budding yeast encodes a single YTH domain-containing protein, PHO92 (<u>Pho</u>sphate metabolism <u>92</u>), that has been originally described to be involved in phosphate metabolism and response (34, 35). Recently, it became clear that PHO92 recognizes m^6^A-modified transcripts, facilitates protein synthesis and subsequent decay of m^6^A-modified transcripts, and promotes meiotic recombination (36, 37). In the fission yeast *Schizosaccharomyces pombe*, an YTH domain-containing protein, Mmi1 (<u>M</u>eiotic <u>m</u>RNA interception 1), was involved in selective elimination of meiosis-specific transcripts via post-transcriptional gene silencing (38). However, Mmi1 lacks the ability to bind to the m^6^A consensus motif, suggesting that the function of YTH domain-containing proteins are not limited to m^6^A recognition and implicated in diverse cellular functions (39). Although m^6^A writers and m^6^A readers have been studied in yeasts, to our best knowledge, m^6^A erasers await its discovery in fungi.

*Fusarium graminearum* is a filamentous phytopathogenic fungus, causing devastating diseases on our staple crops, such as wheat, barley and corn (40). The fungus has served as an excellent model organism for investigating various biological aspects, including host-pathogen interactions, sexual development, mycotoxin production, and RNA editing (41–45). Recently, in the rice blast fungus *Magnaporthe oryzae*, <u>MT</u>-<u>A</u>70 domain protein <u>1</u> (MTA1, orthologous to human METTL4) was shown to play an important role in appressorium formation during infection process via regulation of autophagy process (46, 47). However, the roles of the m^6^A factors are poorly understood in *F. graminearum* and other filamentous fungi. Thus, the aims of this study were to (*i*) investigate phyletic distribution and copy number variations of m^6^A factors in the kingdom Fungi, (*ii*) functionally characterize putative m^6^A factors in *F. graminearum*, and (*iii*) identify potential targets of the m^6^A writer MTA1, which is the sole m^6^A writer found in the Pezizomycotina.

## Results

### Divergence of m^6^A writers in the kingdom Fungi

To resolve phylogenetic relationships of genes encoding an MT-A70 protein domain (Pfam domain: PF05063), we collected 1,568 protein sequences from the UniProt database (accessed on May 21, 2023; https://www.uniprot.org/) and curated the list of MT-A70-containing proteins, excluding redundant entries or sequences with an *E*-value of smaller than 10^-5^ for PF05063 (see Materials and Methods). Following the manual curation, a maximum likelihood tree was reconstructed for 1,116 protein sequences containing an MT-A70 domain in 829 diverse fungal species. This tree revealed the presence of three distinct clades, each including the three previously characterized m^6^A writers in fungi: IME4, KAR4, and MTA1 (Fig. 1A). Within the IME4 and KAR4 clades, there was a branch exhibiting an early divergence, which exclusively consists of sequences derived from early-diverging fungi, such as species belonging to the phyla Chytridiomycota, Mucoromycota, and Zoopagomycota (see clades labelled with red asterisks in Fig. 1A).

**Figure 1.**
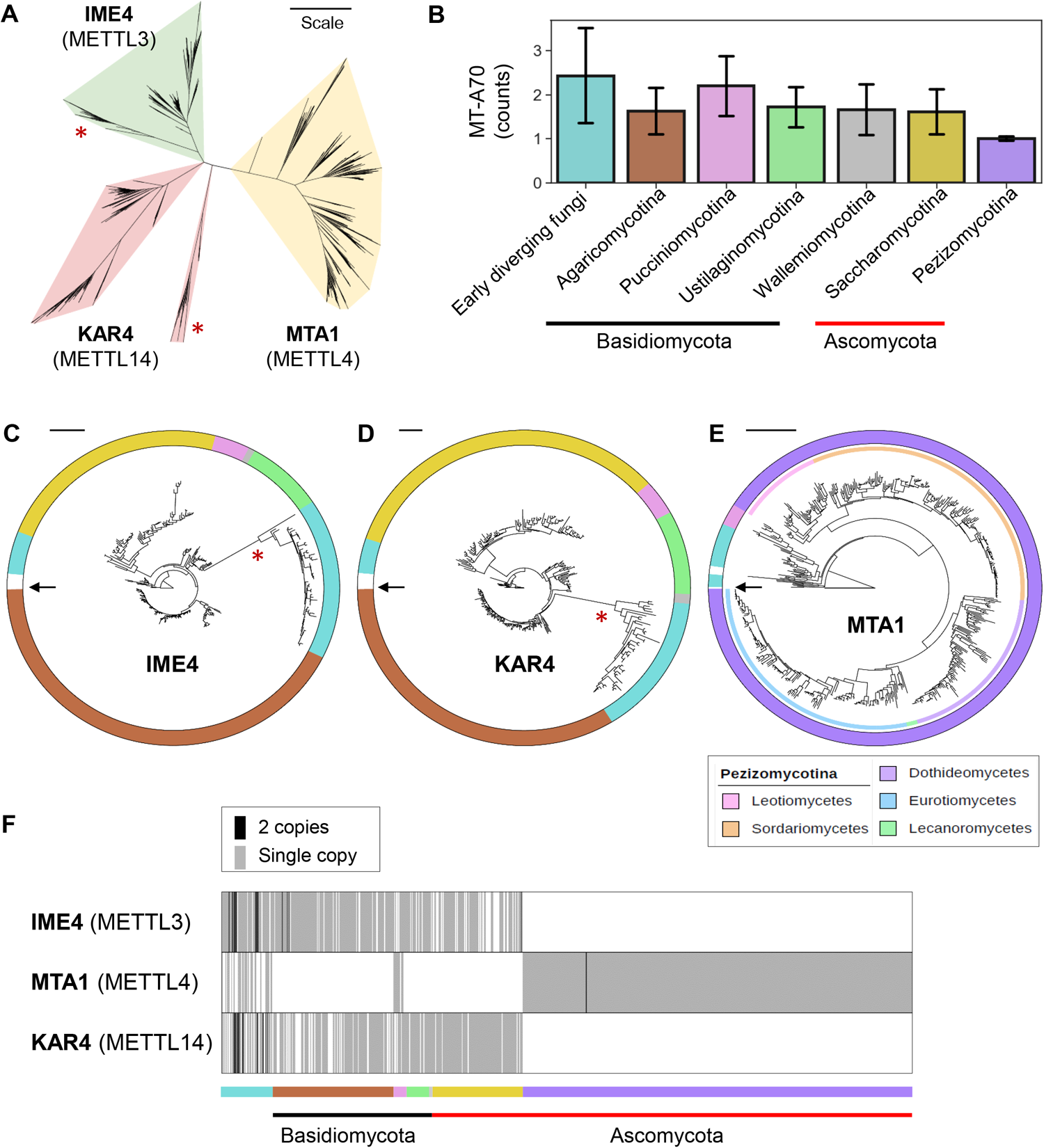
Phylogeny of potential m^6^A writers in fungi. (**A**) The maximum-likelihood phylogeny was estimated from 1,116 sequences of MT-A70 domain-containing proteins identified in 829 fungal species. The tree exhibits three distinct subclades, each corresponding to IME4, KAR4, and MTA1 families. Branches marked with red asterisks suggest clades that likely diverged at earlier time points. The branch lengths in the tree reflect the amount of evolutionary change, with the scale indicating 1.0 amino acid sequence substitution per site. (**B**) Phyletic distribution of MT-A70 domain-containing proteins in fungi. (**C**) The phylogeny was estimated from 308 protein sequences that belong to the IME4 subclade and from five METTL3 orthologues found in metazoans and a model plant *Arabidopsis thaliana*. The arrow indicates *Arabidopsis thaliana* METTL3. The branch marked with a red asterisk suggests clades that likely diverged at earlier time points. The color strip outside the tree represents different fungal taxa, with each color indicating a distinct taxonomic group as shown in Figure 1B. (**D**) The phylogeny was estimated from 298 protein sequences that belong to the KAR4 subclade and from five METTL14 orthologues found in metazoans and a model plant *Arabidopsis thaliana*. The arrow indicates *Arabidopsis thaliana* METTL14. The branch marked with a red asterisk suggests clades that likely diverged at earlier time points. (**E**) The phylogeny was estimated from 510 protein sequences that belong to the MTA1 subclade and from five METTL4 orthologues found in metazoans and a model plant *Arabidopsis thaliana*. The arrow indicates *Arabidopsis thaliana* METTL4. The inner color strip represents fungal class within the subphylum Pezizomycotina, with each color indicating a distinct taxonomic group as shown in the inset box. (**C–E**) More detailed trees showing UniProt protein IDs and bootstrap values can be found in Supplementary Fig. S1. (**F**) Copy number variation of potential m^6^A writers in fungi. The color strip represents different fungal taxa, with each color indicating a distinct taxonomic group as shown in Figure 1B.

Significantly distinct patterns were observed in the phyletic distribution of MT-A70-containing proteins among different taxonomic groups in fungi (Fig. 1B). Notably, many early-diverging fungi and certain species within the Pucciniomycotina subphylum were found to possess more than two MT-A70-containing proteins. The Basidiomycota species (mushroom-forming fungi) and Saccharomycotina species (budding yeasts), typically had one or two MT-A70-containing proteins, whereas nearly all of the Pezizomycotina species that are comprised of entirely of filamentous fungi were found to possess a single MT-A70-containing protein.

Independently reconstructed maximum likelihood trees were generated for the IME4, KAR4, and MTA1 clades including their respective homologs of METTL3, METTL14, and METTL4, which were found in humans (*Homo sapiens*), mouse (*Mus musculus*), zebrafish (*Danio rerio*), fruit fly (*Drosophila melanogaster*), and a model plant (*Arabidopsis thaliana*) serving as an outgroup (Figs. 1C, 1D, and 1E; see Supplementary Fig. S1 for bootstrap values and protein IDs). The phylogenetic patterns of the IME4 and KAR4 clades exhibited resemblance, encompassing early-diverging fungi and species from Basidiomycota and Saccharomycotina. Among species possessing either IME4 (294 species) or KAR4 (286 species), three quarters of the species (221 species) had both m^6^A writers (Supplementary Table S1). As previously mentioned, certain early-diverging fungi exhibited the presence of two copies of IME4 and KAR4. One of these copies appears to have undergone divergence before the emergence of Dikarya (commonly known as “higher fungi”), forming distinct clades with robust 100% bootstrap support (Fig. 1F; Supplementary Fig. S1). The phylogeny of MTA1 exhibited significant monophyly and demonstrated topological congruence with the species phylogeny of Pezizomycotina (Fig. 1E). While species belonging to Pezizomycotina possessed MTA1 as a sole MT-A70-containing protein, many early diverging fungi and species from Pucciniomycotina exhibited the presence of three m^6^A writers, IME4, KAR4, and MTA1 (Fig. 1F).

Recent studies revealed the components of yeast m^6^A methyltransferase complexes were highly conserved with those found in mammals, insects, and plants, including IME4, KAR4, MUM2 (a homolog of WTAP), and VIR1 (32, 33). Thus, we conducted a search in the UniProt database for relevant Pfam domains, specifically PF17098 for the WTAP/Mum2p family and PF15912 for the N-terminal domain of *virilizer* (hereafter coined VIRN). We identified 172 and 53 fungal proteins that harbor PF17098 for WTAP and PF15912 for VIRN, respectively, in the current UniProt database (accessed on June 4, 2023; Supplementary Table S1). After manual curation, we reconstructed phylogenetic trees for WTAP and VIRN, together with homologs found in humans, mouse, zebrafish, fruit fly, and *Arabidopsis thaliana* as an outgroup. WTAP orthologs including Mum2 in yeasts were found only in early-diverging fungi and species from Basidiomycota, and Saccharomycotina, lacking in Pezizomycotina species (Supplementary Fig. S1). Only 37 fungal species that belong to early-diverging fungi and Basidiomycota were shown to have genes encoding a VIRN domain (Supplementary Fig. S1). The paucity of homologous sequences in fungi may be attributable to the fact that WTAP and VIRN sequences have diverged significantly from the known sequences, making their identification challenging especially in Ascomycetous fungi, such as Saccharomycotina and Pezizomycotina. Indeed, a homolog of the virilizer protein, VIR1, was recently found in the budding yeast, which lacks the apparent Pfam domain for VIRN, but it exhibits a 3D structure similar to the human homolog VIRMA (32).

### Identification of putative m^6^A reader and eraser in Fungi

Among RNA-binding proteins specifically recognizing and binding to m^6^A, the best studied readers are YTH domain-containing proteins. PHO92 was found as the only protein containing the YTH domain in the budding yeast, while animals can have up to five such proteins and plants can have more than 10. A recent study revealed target transcripts of PHO92 and its important roles during meiosis in the budding yeast (37). However, little is known about m^6^A readers in other fungal taxa. Therefore, we searched for proteins containing an YTH domain (PF04146) in the UniProt database (accessed on May 21, 2023) and found 2,029 fungal sequences. To shorten the list of potential m^6^A readers, entries with irrelevant protein names (*e.g.* DNA repair protein RAD51) were excluded, and protein sequences below 300 or above 1,000 amino acids in length were filtered out. After manual curation, we reconstructed a phylogenetic tree for the final list of 868 YTH domain-containing proteins in 695 diverse fungal species (Fig. 2A; see Supplementary Fig. S1 for bootstrap values and protein IDs). The tree was rooted to YTH domain-containing proteins outside of fungi, including human YTHDF2. The tree was composed of three main clades that were strongly supported by high bootstrap values. One clade contained PHO92 homologs from species belonging to Saccharomycotina and Basidiomycota, as well as YTH domain-containing proteins found in mammals, an insect, and a plant. Another clade exclusively comprised species from Pezizomycotina and was placed sister to the PHO92 clade. The third clade, here dubbed YTH1, consisted of early-diverging fungi and species from Basidiomycota and Ascomycota, displaying a topology that largely matched the species phylogeny. Approximately half of the Agaricomycotina species (67 out of 128) and one fifth of the Pezizomycotina species (98 out of 498) examined in this study were found to possess both PHO92 and YTH1, while early-diverging fungi and species belonging to Pucciniomycotina, Ustilaginomycotina, and Wallemiomycotina, were found to have only PHO92 (Figs. 2B and 2C). While the majority of species belonging to Saccharomycotina (30 out of 37) were found to possess PHO92, there were several species including *Yarrowia lipolytica* (a lipid-producing yeast) that lacked PHO92 and instead exhibited the presence of YTH1 (Fig. 2C; Supplementary Table S1).

**Figure 2.**
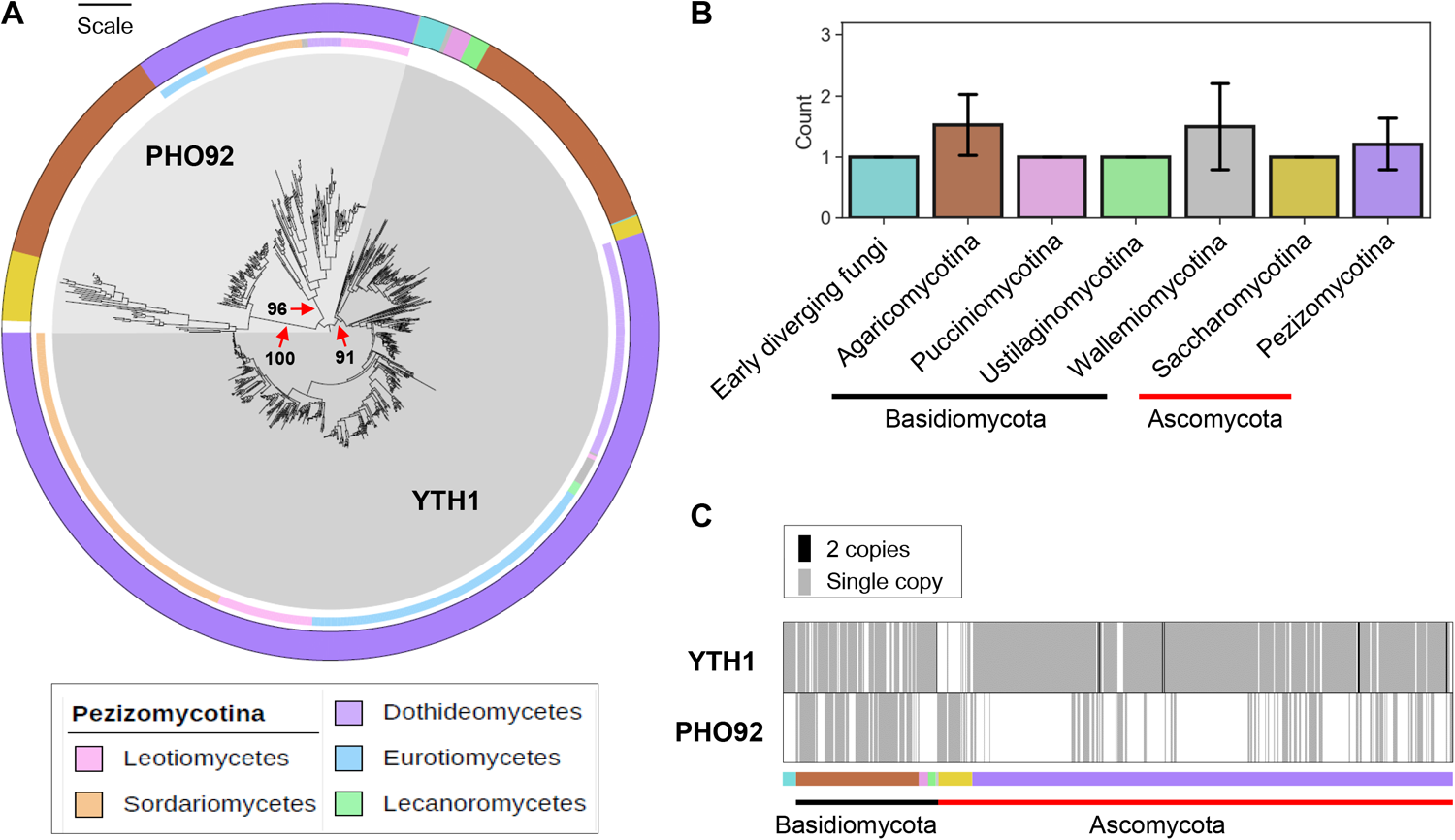
Phylogeny of potential m^6^A readers in fungi. (**A**) The maximum-likelihood phylogeny was estimated from 868 sequences of YTH domain-containing proteins identified in 695 fungal species. Red arrows indicate highly supported clades representing PHO92 (Agaricomycotina and Saccharomycotina), PHO92 (Pezizomycotina) and YTH1 families. The branch lengths in the tree reflect the amount of evolutionary change, with the scale indicating 1.0 amino acid sequence substitution per site. The outer color strip represents subphyla, with each color indicating a distinct taxonomic group as shown in (B), and the inner color strip represents fungal class within the subphylum Pezizomycotina, with each color indicating a distinct taxonomic group as shown in the inset box. (**B**) Phyletic distribution of YTH domain-containing proteins in fungi. (**C**) Copy number variation of potential m^6^A readers in fungi. The color strip represents different fungal taxa, with each color indicating a distinct taxonomic group as shown in Figure 2B.

2OG-Fe[II] dioxygenase AlkB-like domain-containing proteins were known to play roles as m^6^A erasers by removing the methyl group from m^6^A in mammals and plants (26, 48–50). However, there has been no information on AlkB homologs (ALKBHs) in fungi. Thus, we searched for protein sequences containing a 2OG-Fe(II) dioxygenase superfamily domain in the genome of *F. graminearum* strain PH-1. We identified five genes harboring a 2OG-Fe(II) dioxygenase domain (PF13532) with an *E*-value of smaller than 10^-10^ (gene ID: FGRRES_01255, FGRRES_09872, FGRRES_16456, FGRRES_16652, and FGRRES_20373) (Table 1). 2OG-Fe(II) dioxygenases are involved in diverse cellular processes, such as demethylation of DNA/RNA and modification of histone proteins, as well as secondary metabolite production. To get insights into evolutionary relationships of different types of ALKBH, we reconstructed a phylogenetic tree of ALKBH proteins found in 27 selected fungi, including several functionally characterized ALKBH proteins in humans and Arabidopsis (Fig. 3). Early-diverging fungi, such as *Basidiobolus meristosporus* and *Spizellomyces punctatus*, possessed 10 and 9 2OG-Fe(II) dioxygenases, whereas yeasts that belong to Saccharomycotina tend to have only one or two (Supplementary Table S2). ALKBH2 (FGRRES_09872), ALKBH3 (FGRRES_16456), and ALKBH4 (FGRRES_01255 were closely-related to human ALKBH3 and Arabidopsis ALKBH2, which are responsible for repairing DNA damage caused by alkylation (51, 52). ALKBH5 (FGRRES_20373) was placed sister to the clade containing human ALKBH6, which plays a role in DNA repair (53, 54). ALKBH1 (FGRRES_16652) formed a well-supported clade with human and Arabidopsis ALKBH1s, known for their versatile functions in removing methyl groups from DNA/RNA, as well as modifying histone H2A (55–57). It was shown that human ALKBH5 and Arabidopsis ALKBH9B and ALKBH10B demethylate m^6^A (48, 58–60). These m^6^A erasers in mammals and plants formed a clade ancestral to clades including ALKBH1 and ALKBH5 in *F. graminearum* (Fig. 3).

**Figure 3.**
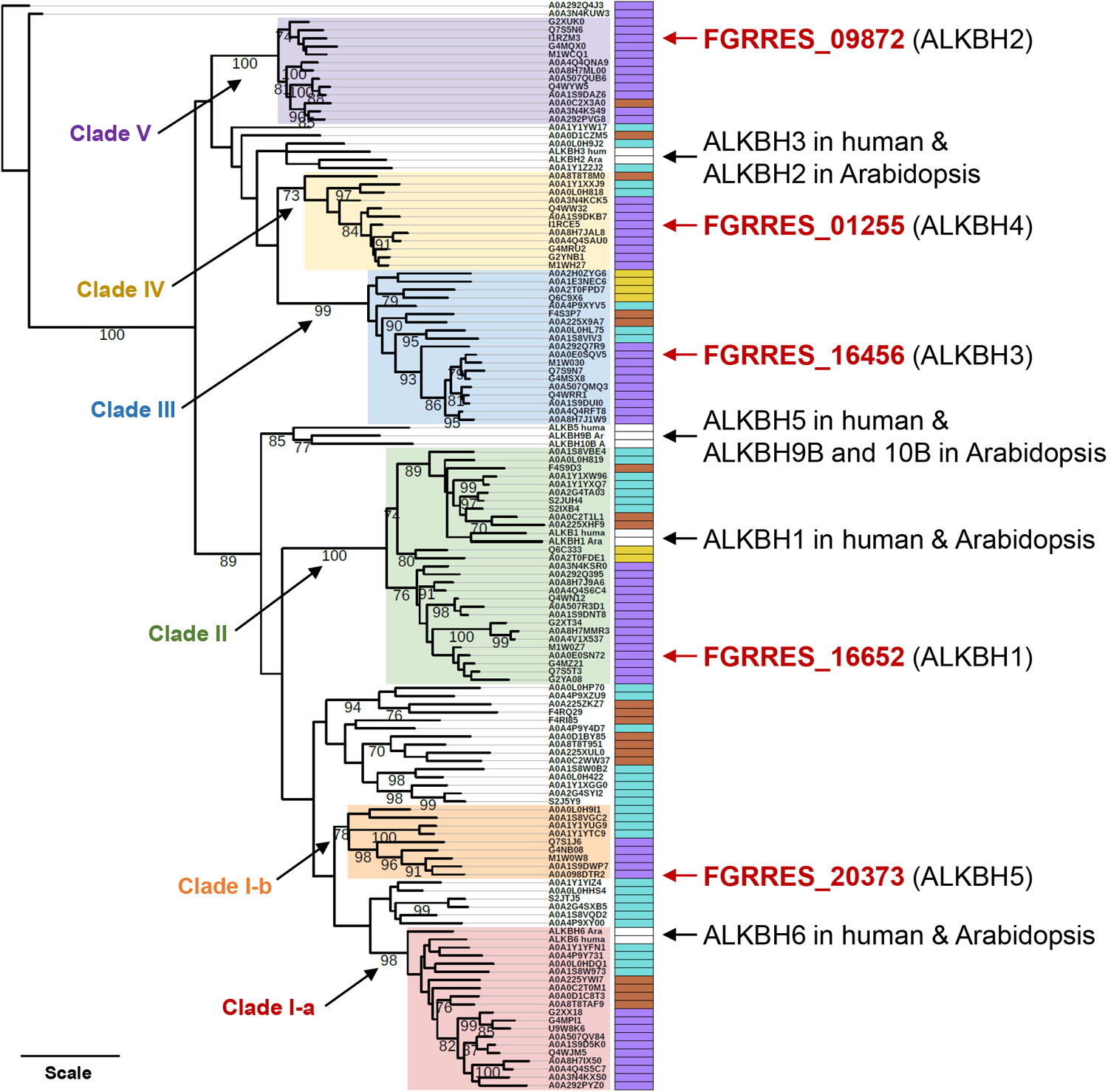
Phylogeny of potential m^6^A erasers in fungi. The maximum-likelihood phylogeny was estimated from 125 sequences of 2OG-Fe(II) dioxygenase AlkB-like domain-containing proteins identified in 27 selected fungal species. The color strips on the right side of leaves indicate different fungal taxa: purple**–**Pezizomycotina, yellow**–** Saccharomycotina, blown**–**Basidiomycota, cyan**–**early-diverging fungi, white**–**human or *Arabidopsis thaliana*. Highly supported clades each containing an ALKBH identified in *F. graminearum* were shaded with different colors. Bootstrap values of greater than 70% were shown. Branch lengths are proportional to the inferred amount of evolutionary change, and the scale represents 1.0 amino acid sequence substitutions per site.

**Table 1.** Putative m^6^A factors found in *F. graminearum*.

| **Name** | **ID**<br>**(FGRRES)** | **AA** | **Pfam** | **E-value** | **Putative**<br>**function** |
| --- | --- | --- | --- | --- | --- |
| MTA1 | 06225 | 333 | PF05063 | 1.2e-34 | writer |
| WTAP<sup>1</sup> | 01626 | 266 | PF17098 | 0.069 | writer |
| VIRN<sup>1</sup> | 06249 | 143 | PF15912 | 0.073 | writer |
| YTH1 | 01159 | 613 | PF04146 | 9.9e-66 | reader |
| ALKBH1 | 16652 | 346 | PF13532 | 3.5e-30 | eraser |
| ALKBH2 | 09872 | 933 | PF13532 | 1e-29 | eraser |
| ALKBH3 | 16456 | 443 | PF13532 | 5.6e-35 | eraser |
| ALKBH4 | 01255 | 326 | PF13532 | 2.8e-28 | eraser |
| ALKBH5 | 20373 | 228 | PF13532 | 6.2e-15 | eraser |

<sup>1</sup> E-value is greater than 10<sup>-5</sup>

### Functional characterization of putative m^6^A factors in *F. graminearum*

In order to examine the functions of m^6^A factors in *F. graminearum* during sexual development, we conducted a search for Pfam domains associated with m^6^A writer, reader, and eraser, and discovered several potential m^6^A factors (Table 1). Among them, *F. graminearum* possessed genes homologous to *MTA1* and *YTH1*, along with the previously mentioned five *ALKBHs*. Additionally, we identified two genes that encoded hypothetical proteins harboring PF17098 for WTAP and PF15912 for VIRN, respectively, with *E*-values of greater than 10^-5^ (Table 1). Despite the low sequence similarity to *WTAP* and *VIRN* found in fungi, we included these potential m^6^A writers in the knockout study because their expression levels were observed to increase during sexual development (see Fig. 6 in the section “Roles of MTA1 in conidial germination and hyphal growth”). We individually deleted m^6^A factors in *F. graminearum* strain PH-1 (wild-type, WT). The authenticity of the resulting knockout mutants were verified by diagnostic PCRs (Fig. 4A). The knockouts displayed normal growth and produced perithecia, which are characterized by their dark purple coloration and flask-shaped or spherical structures (Fig. 4B). Morphologies of ascospores (the products of meiosis) enveloped in sac-like structures called asci were also normal in the knockouts, indicating that m^6^A factors did not affect sexual development in *F. graminearum* (Supplementary Fig. S2).

**Figure 4.**
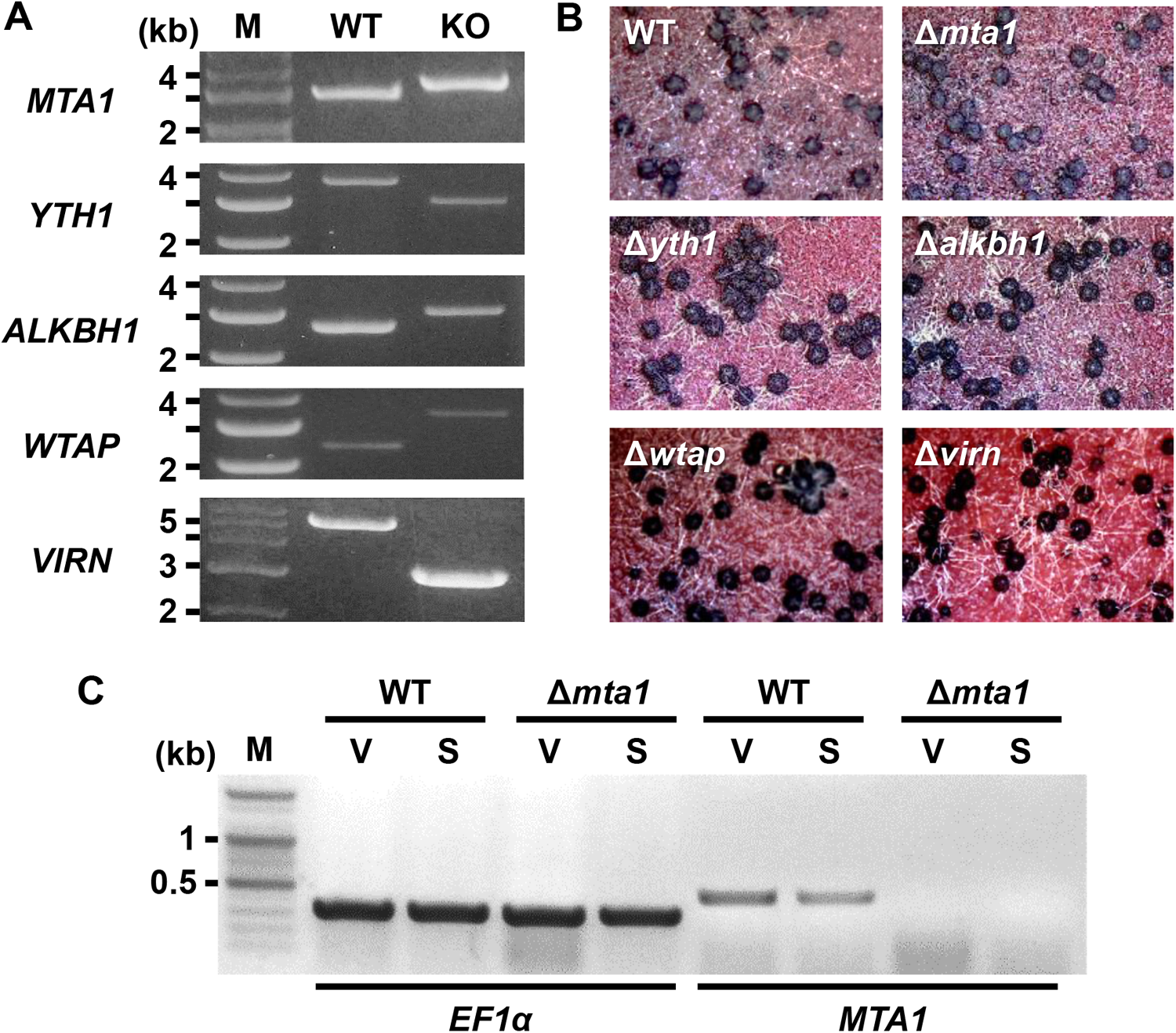
Generation of knockout mutants lacking potential m^6^A factors in *F. graminearum*. (**A**) Diagnostic PCRs confirmed homologous integration of the split marker constructs to the target gene loci. Note the PCR band size between the wild-type (WT) and knockout strains (KO) due to the gene replacement. M–1-kb DNA ladder. (**B**) Perithecia production of the WT and knockout strains grown on carrot agar media. Photographs were taken with a dissecting microscope at 6 days after induction of sexual development. (**C**) RT-PCR analyses of the WT and Δ*mta1* strains. Since *MTA1* lacks introns, distinguishing RNA expression from possible genomic DNA contamination in RT-PCR analysis was challenging. To verify RNA integrity, a primer set was designed for amplifying *EF1α*, including two introns. Notably, no band corresponding to genomic DNA amplicon for the EF1α reference gene was observed (658 bp for gDNA, 356 bp for mRNA), indicating the absence of genomic DNA contamination. Expression of *MTA1* was confirmed in the WT, whereas no discernible band was observed in Δ*mta1* strain. The samples labeled V were collected before sexual induction (*i.e.* vegetative growth stage), samples labeled S were collected six days after sexual induction (*i.e.* sexual growth stage). M–100-bp DNA ladder.

Since m^6^A writer, *IME4,* in the budding yeast plays a crucial role in initiating sexual development, we carefully examined the genotype of Δ*mta1*. Although genotyping confirmed the deletion of *MTA1* in Δ*mta1*, we were intrigued to find that a PCR band specific to *MTA1* could still be amplified from Δ*mta1*. This suggested that Δ*mta1* is heterokaryotic, indicating that it still contains one or more “wild-type” nuclei. To obtain homokaryons, we performed additional single spore isolations, using Δ*mta1*, and examined 10 single-spored isolates derived from Δ*mta1*. Diagnostic PCR analysis confirmed the deletion of *MTA1* in all 10 isolates, yet a fragment of *MTA1* was still amplified from these isolates (Supplementary Fig. S3). The expression levels of *MTA1* were markedly reduced in Δ*mta1*, suggesting a lower abundance of wild-type nuclei compared to the WT (Fig. 4C). This observation was further supported by RNA-seq analysis. When examining the mapped reads on the *MTA1* locus in both WT and Δ*mta1*, we found a decrease in the number of reads mapped to the *MTA1* locus in Δ*mta1* compared to the WT (Supplementary Fig. S3). The presence of RNA-seq reads mapping to the *MTA1* locus, which should have been absent in homokaryotic Δ*mta1*, served as additional proof for the heterokaryotic nature of Δ*mta1* and indicated that Δ*mta1* is, in fact, a knockdown strain. Consequently, we will refer to Δ*mta1* as the *MTA1*-KD1 strain hereafter.

### MTA1 as an m^6^A writer in *F. graminearum*

Extensive efforts were unsuccessful to obtain knockouts completely lacking *MTA1* (Supplementary Fig. S3), presumably due to its indispensable role in *F. graminearum*. To get better insight into the roles of m^6^A RNA methylation in *F. graminearum*, we generated strains overexpressing *MTA1*. RT-PCR analysis confirmed that two strains, *MTA1*-OE4 and *MTA1*-OE5, exhibited overexpression of *MTA1* (Fig. 5A). It was notable that *MTA1* was overexpressed to a greater extent in the *MTA1*-OE5 strain, compared to *MTA1*-OE4. Both semi-quantitative RT-PCR and RNA-seq analyses indicated that the expression level of the *MTA1*-OE5 strain was approximately 100 times greater than that of the WT (Supplementary Fig. S4). The WT and *MTA1*-OE4 strains produced normal asci containing mature ascospores six days after the induction of sexual development, while it was not until 11 days after the induction that the *MTA1*-OE5 strain produced fully developed asci (Fig. 5B). The difference in the observed phenotypes in the two *MTA1*-overexpressing strains could likely be attributable to the variation in the degrees of *MTA1* expression, and thus the *MTA1*-OE5 strain which displayed substantial overexpression of *MTA1* was used for further analysis and investigation. To determine whether MTA1 is required for m^6^A RNA methylation in *F. graminearum*, we compared total m^6^A RNA methylation levels between the WT and *MTA1*-OE5 strain. The amount of m^6^A RNA in WT was 0.075 ± 0.015% of the total RNA, which was approximately half of that in the *MTA1*-OE5 strain (0.152 + 0.033% of the total RNA), which indicated that the m^6^A RNA methylation level was significantly increased in the *MTA1*-OE5 strain compared to the WT (Fig. 5C).

**Figure 5.**
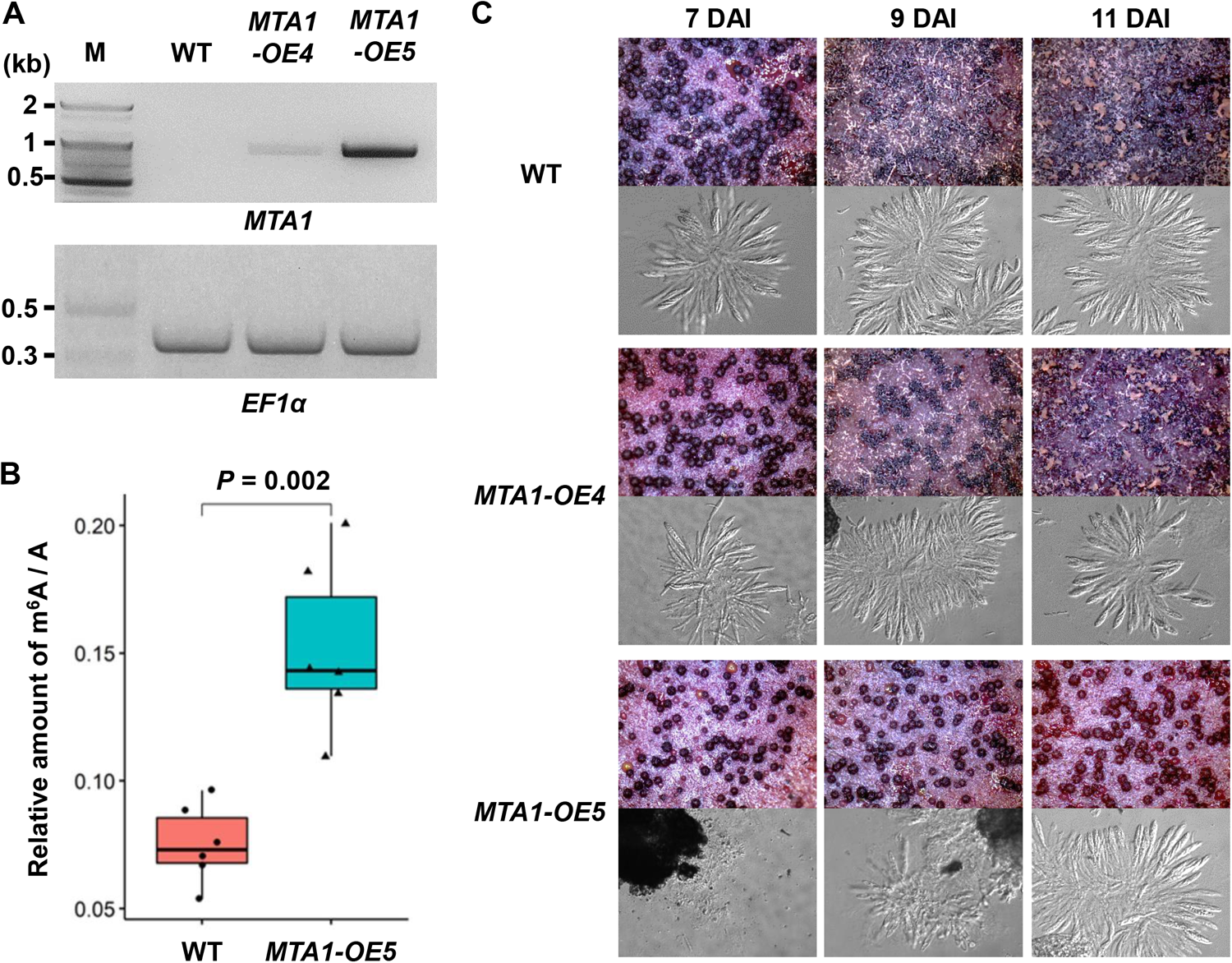
Overexpression of the m^6^A writer *MTA1*. (**A**) RT-PCR analyses of the WT and *MTA1*-overexpressing strains (*MTA1*-OE4 and *MTA1*-OE5). Since *MTA1* lacks introns, distinguishing RNA expression from possible genomic DNA contamination in RT-PCR analysis was challenging. To verify RNA integrity, a primer set was designed for amplifying *EF1α*, including two introns. Notably, no band corresponding to genomic DNA amplicon for the EF1α reference gene was observed (658 bp for gDNA, 356 bp for mRNA), indicating the absence of genomic DNA contamination. Overexpression of *MTA1* was confirmed in the *MTA1*-OE4 and *MTA1*-OE5 strains. Note that *MTA1* was significantly overexpressed in the *MTA1*-OE5 strain, compared to the WT, from which no discernible band was observed at PCR cycle of 30. M–100-bp DNA ladder. (**B**) m^6^A methylation level in RNA extracted from the WT and *MTA1*-OE5 strains. Mann-Whitney U test was performed to compare the means of the ratio for m^6^A to A between the WT and *MTA1*-OE5 strains. Box and whisker plots indicate the median, interquartile range between the 25th and 75th percentiles (box), and 1.5 interquartile range (whisker). (**C**) Perithecia production of the WT, *MTA1*-OE4 and *MTA1*-OE5 strains grown on carrot agar media (upper panels). Photographs were taken with a dissecting microscope at the indicated days after induction of sexual development (DAI). Squash mounts of perithecia were observed with a compound microscope (lower panels, 400**×** magnification). Mature ascospores were observed in the WT and *MTA1*-OE4 strains as early as 7 DAI, whereas in the *MTA1*-OE5 strain, it was not until 11 DAI that ascospore formation became evident.

### Roles of *MTA1* in conidial germination and hyphal growth

In previous studies, we obtained transcriptome data for conidial germination on Bird agar medium (61), as well as for perithecial development on Carrot agar medium (62). By integrating these datasets with the RNA-seq data obtained in the present study (freshly-harvested conidia), we conducted a comprehensive analysis of expression level changes in m^6^A factors throughout the life cycle of *F. graminearum*, encompassing various stages of vegetative growth and perithecial development (Fig. 6A). The m^6^A writer *MTA1* and a potential m^6^A reader *YTH1* demonstrated a synchronized expression pattern, as indicated by a high Pearson correlation coefficient of 0.94. Their expression levels exhibited a significant increase during stage 1 (S1, 15 minutes after incubation of conidia on Bird agar medium), followed by a decline during hyphal growth (S2 and S3), and remained at a basal level during perithecial development (S4–S9). Interestingly, the expression levels of *ALKBH1* and *ALKBH5*, the two most probable m^6^A erasers in *F. graminearum*, reached their peak at stage 2 (S2, 3 h after incubation), at which conidia germinated and hyphae started to extend (Supplementary Fig. S5). During active hyphal growth at stage 3 (S3, 11 h after incubation), the major components of fungal cytoskeleton, actin (FGRRES_07735) *α*-tubulin (FGRRES_00639), and *β*-tubulin (FGRRES_09530) that play a crucial role in polarity establishment, maintenance and polar growth, displayed the greatest induction during the life cyle of *F. graminearum*.

**Figure 6.**
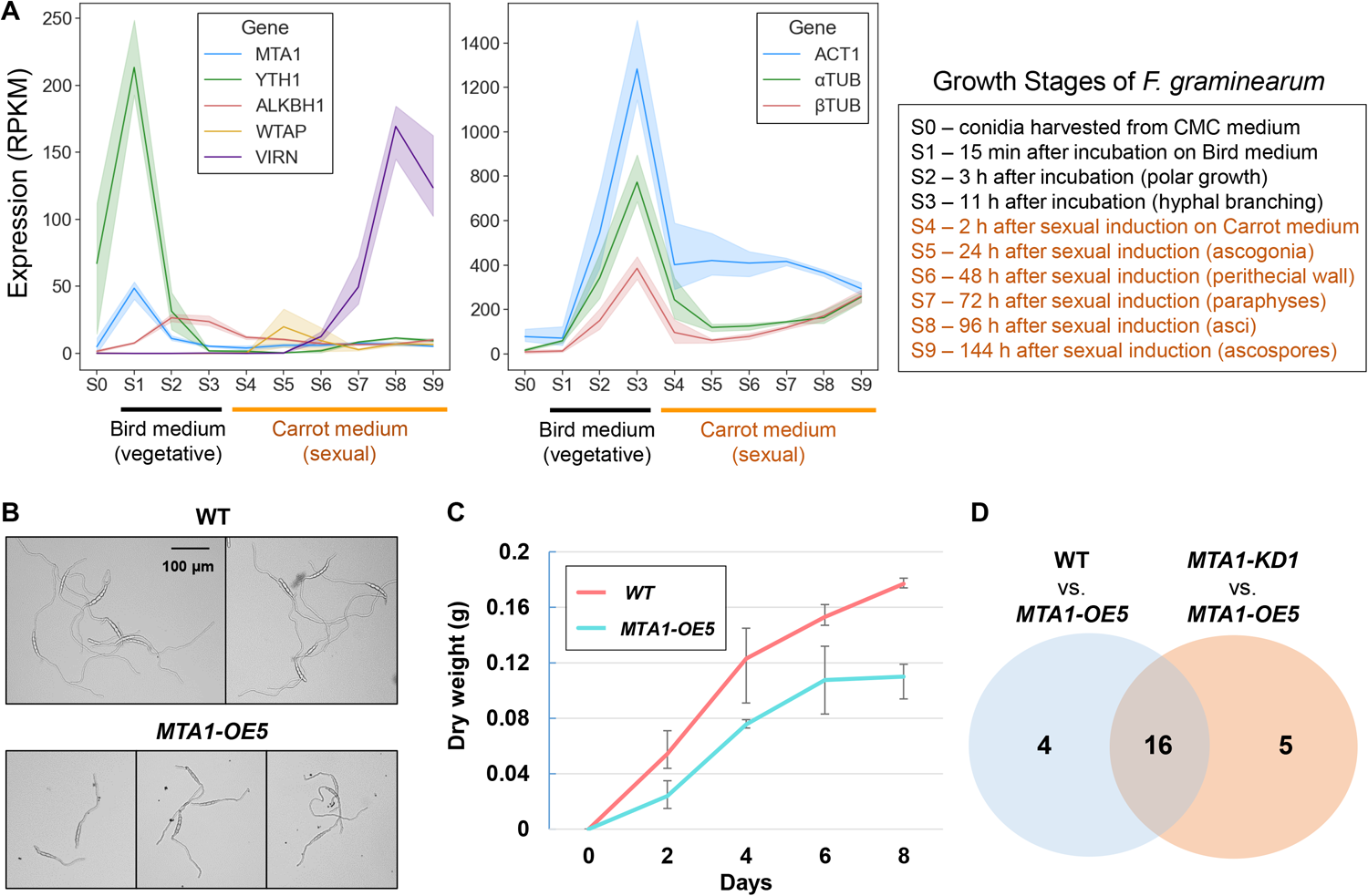
Conidial germination affected by the m^6^A writer *MTA1*. (**A**) Gene expression profiles of potential m^6^A factors (left panel) and housekeeping genes (right panel) in *Fusarium graminearum*. Average values for reads per kilobase per million mapped reads (RPKM) values for three replicates samples were plotted. Bands surrounding the line plots indicate 95% confidence intervals of the means. The *x*-axis are different growth stages of *F. graminearum* (S0–S9). See the right box for the description of vegetative and sexual growth stages. Gene ID for potential m^6^A factors are ALKBH1 (FGRRES_16652), MTA1 (FGRRES_06225), VIRN (FGRRES_06249), and WTAP (FGRRES_01626). Housekeeping genes examined here are *α*-tubulin (*α*TUB, FGRRES_00639), *β*-tubulin (*β* TUB, FGRRES_09530), and actin 1 (ACT1, FGRRES_07735). (**B**) Conidia germination and polar growth of hyphae in quarter-strength potato dextrose broth (q-PDB) medium. Photos were taken 16 hours after incubation. Note that shorter hyphae germinated from macroconidia of the *MTA1*-OE5 strain, compared to the wild-type (WT) strain. (**C**) Dry weight of mycelia grown in q-PDB medium was measured with two days interval. (**D**) The numbers of differentially expressed genes in fresh macroconidia harvested from carboxymethylcellulos (CMC) medium between the WT and *MTA1*-OE5 strains, and between the *MTA1*-KD1 and *MTA1*-OE5 strains.

Taking into account the expression profile, we proceeded to investigate the phenotypic characteristics of the *MTA1*-OE5 strain during conidial germination and hyphal growth. In contrast to the WT, the hyphal tips did not exhibit elongation in an overnight culture of the *MTA1*-OE5 strain cultivated in quarter-strength potato dextrose broth (q-PDB) medium (Fig. 6B). Furthermore, in the *MTA1*-OE5 strain growing on potato dextrose agar (PDA) medium, the hyphae at the leading edge displayed a tendency to have fewer branches compared to the WT (Supplementary Fig. S6). Next, we conducted measurements of the total biomass of the WT and *MTA1*-OE5 strains cultured in q-PDB medium for a duration of 8 days. The *MTA1*-OE5 strain exhibited a slower growth rate compared to the WT, particularly during the initial two days of cultivation (Fig. 6C). To investigate genes responsible for the delayed conidial germination observed in the *MTA1*-OE5 strain, we performed differential gene expression analyses between the WT, *MTA1*-KD1, and *MTA1*-OE5 strains, using RNA-seq data obtained from freshly harvested conidia (stage 0, S0). In this analysis, we identified a total of 20 differentially-expressed (DE) genes (|log2 fold-change| > 2), when comparing the WT and *MTA1*-OE5 strains, as well as 21 DE genes when comparing the *MTA1*-KD1 and *MTA1*-OE5 strains (Fig. 6D). Notably, 16 genes were found to be commonly differentially expressed in both comparisons, including *MTA1*.

There was no DE genes observed between the WT and *MTA1*-KD1 strains (Supplementary Fig. S7). Among the DE genes, we found that 17 genes were transcriptionally upregulated, and 8 genes were downregulated in the *MTA1*-OE5 strain (Table 2). The DE genes included 9 genes encoding hypothetical proteins without any predicted protein domain and 16 functioanlly annotated genes encoding two transcription factors, two transporters, and diverse enzymes.

**Table 2.**
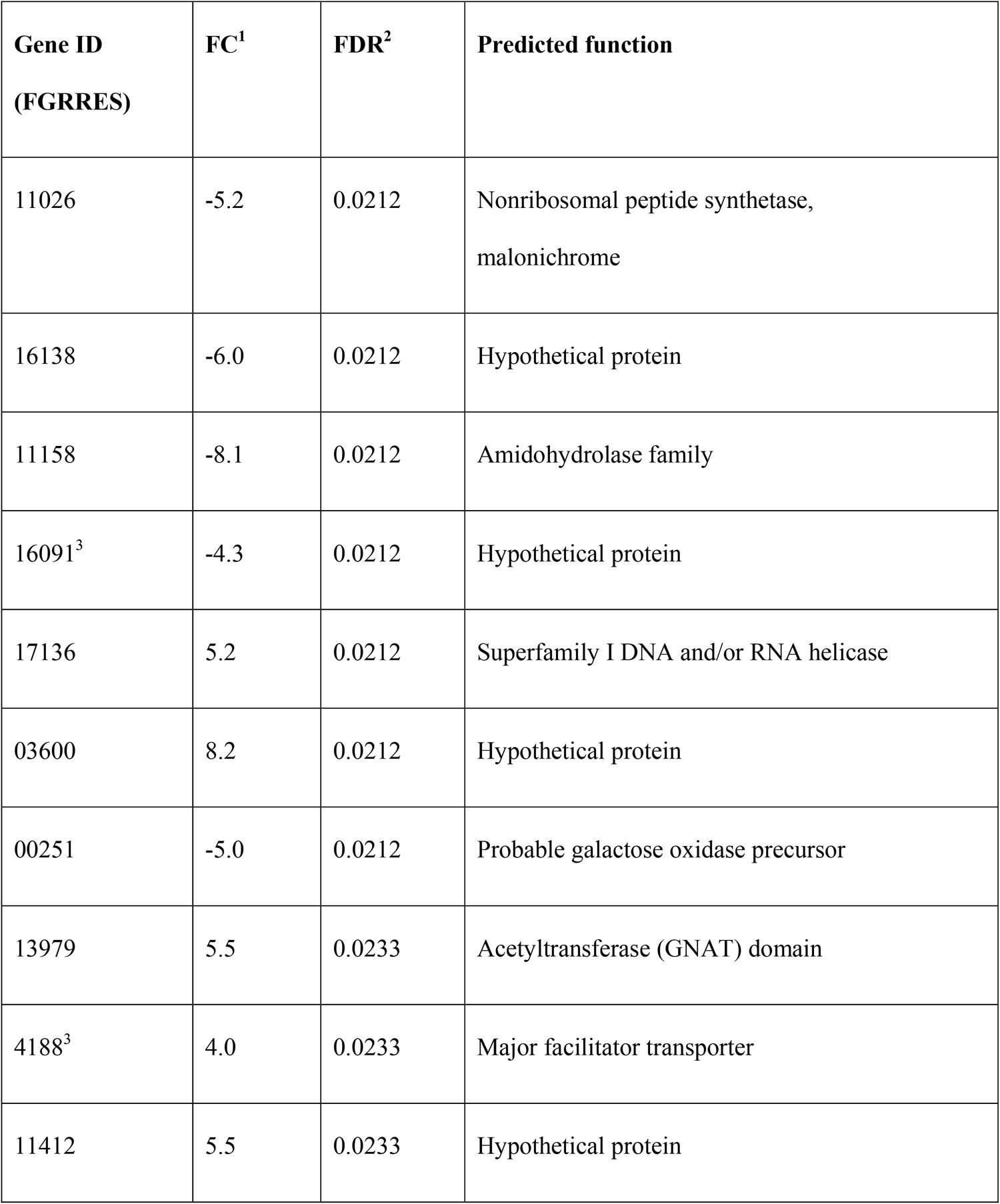
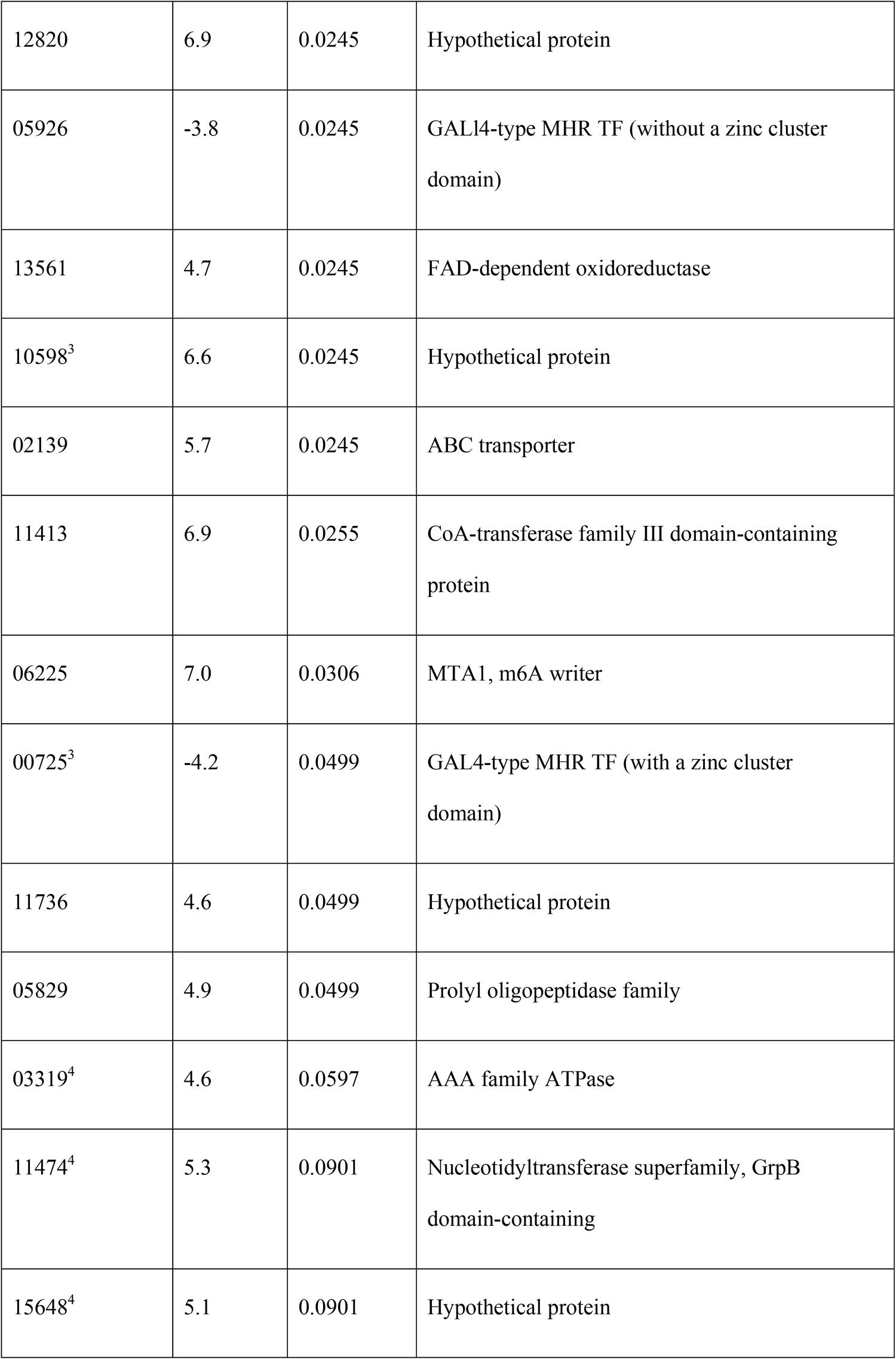
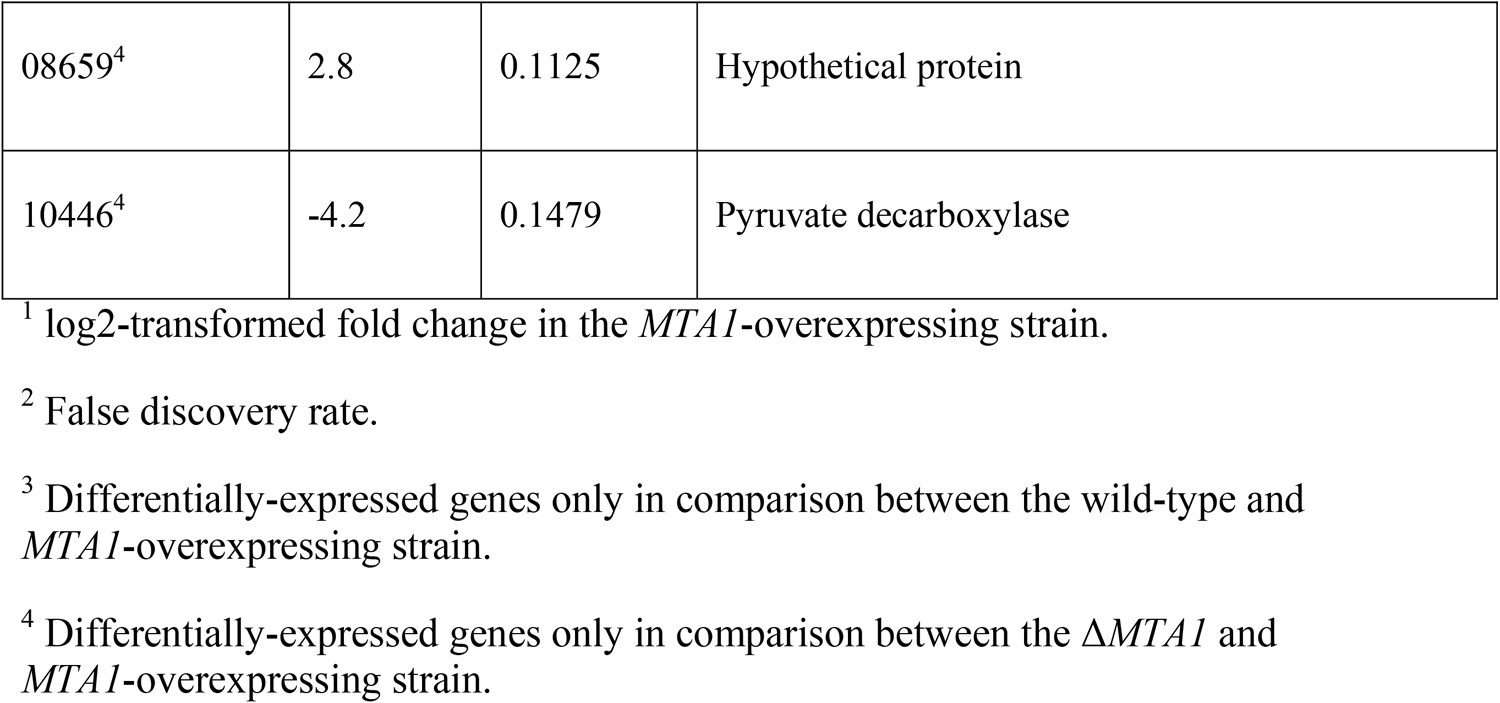
Differentially expressed genes in the *MTA1*-overexpressing strain.

## Discussion

### Roles of m^6^A writer in filamentous fungi

In the phylum Ascomycota, the biological functions of m^6^A writers in budding yeasts (Saccharomycotina) and filamentous fungi (Pezizomycotina) may have evolved independently. Species belonging to the Pezizomycotina possess only single m^6^A writer *MTA1*, a homolog of *METTL4*, while species belonging to the Saccharomycotina have two m^6^A writers, *IME4* and *KAR4* that are homologs of *METTL3* and *METTL14*, respectively.

In the budding yeast, *IME4* and *KAR4* are crucial for initiation of meiosis for the ascospore formation (29, 63). However, the m^6^A writer *MTA1* is dispensable for perithecial development and ascospore production in filamentous fungi, *F. graminearum* and *M. oryzae* (46). The relative abundance of m^6^A in *M. oryzae* (0.069 ± 0.003%) was comparable to that in *F. graminearum* (0.075 ± 0.015%) (47), suggesting that the degree of m^6^A levels of total mRNA is similar in these two filamentous fungi. In the rice blast fungus *M. oryzae*, a knockout strain lacking *MTA1* showed a defect in appressorium formation during the infection process on epidermal cells of rice (47). Expression levels of autophagy-related genes in *M. oryzae* were changed according to the degree of m^6^A level, which is important for normal appressorium formation. Although the homozygous Δ*mta1* strain in *M. oryzae* was viable, our extensive efforts to obtain homozygous Δ*mta1* was unsuccessful in *F. graminearum*, suggesting its essential role. Given the significant impact on conidial germination and hyphal growth observed in the *MTA1*-overexpressing strain in *F. graminearum*, the role of *MTA1* appears to have diverged in the two plant pathogenic fungi.

Most importantly, *F. graminearum* does not form appressoria and utilizes distinct strategies to penetrate epidermal cells of the host plants. Both *F. graminearum* and *M. oryzae* belong to the Sordariomycetes, but fall into different orders, Hypocreales and Magnaporthales, respectively (64). RNA-seq analyses indicated significant divergence in gene expression including autophagy-related genes during conidial germination and infection processes between *F. graminearum* and *M. oryzae* (61). Although we did not assess the virulence of the *MTA1*-OE5 strain due to their abnormal growth phenotype, it is most likely that the *MTA1*-OE5 strain would exhibit reduced virulence on host plants, compared to the WT. The m^6^A writers *IME4* and *KAR4* were crucial for the initiation of sexual cycle and formation of ascospores in the budding yeast. However, the m^6^A writers *MTA1* does not appear to be responsible for perithecial development and ascospore production in *F. graminearum* and *M. oryzae*. Although the *MTA1*-OE5 strain exhibited delayed perithecial development, it produced normal ascospores. In the Pezizomycotina, A-to-I RNA editing, one of the eukaryotic RNA modifications, is crucial for sexual development and ascospore formation (43–45, 65–67).

### Evolutionary perspectives of m^6^A factors in fungi

Several ancestral traits observed in early-diverging fungi are shared with metazoans or unicellular opisthokonts, which have been subjected to extensive parallel loss across the Dikarya lineages (68, 69). As with metazoans, many species of early-diverging fungi and the Pucciniomycotina (the earliest-diverging subphylum within Basidiomycota) possess all three m^6^A writers, IME4, MTA1, and KAR4. These suggested that budding yeasts (Saccharomycotina) may have lost the m^6^A writer MTA1. Conversely, species belonging to Pezizomycotina, which entirely consist of filamentous fungi, may have lost the m^6^A writers IME4 and KAR4. The phyletic distribution of m^6^A writers in fungi suggested that Pezizomycotina species including *F. graminearum* and *M. oryzae* have likely embraced m^6^A machineries, such as *MTA1*, as a means to regulate filamentous growth and facilitate host penetration. This adoption of m^6^A machinery may have played a crucial role in the development and maintenance of the filamentous morphology that is characteristic of these fungi, allowing them to effectively colonize and interact with their respective hosts.

In contrast to well-characterized yeast m^6^A methyltransferase complexes including *IME4* and *KAR4* m^6^A writers, our understanding of m^6^A machineries in filamentous fungi is still limited. Despite our functional characterization of potential *WTAP* and *VIR1* homologs in *F. graminearum*, it is unlikely that these represent the genuine homologs. This is because the homologs of *WTAP*/*MUM2* and *VIR1* are missing in species within all the Pezizomycotina species. The failure to identify homologs of *WTAP* and *VIR1* in our study is likely due to significant sequence divergence of these m^6^A factors within the Pezizomycotina. The sequence differences might have hindered their detection using conventional homology search methods, indicating that these m^6^A factors in Pezizomycotina may have undergone substantial changes in the primary amino acid sequences. Until recently, the existence and function of *VIR1* in the budding yeast had remained unknown, but through the application of protein folding prediction tools, its identity was revealed (32). Recent advancements in protein 3D structure prediction tools to uncover hidden homologs (70, 71) hold promise for identifying homologous members of the m^6^A methyltransferase complexes in Pezizomycotina species. Alternatively, it is possible that components of m^6^A machineries associated with MTA1 are completely distinct from those found in the yeast m^6^A methyltransferase complexe associated with IME4 and KAR4. In this case, it will be necessary to identify components that interact with MTA1 via co-immunoprecipitation.

In addition, it is worth to mention that, considering the number of putative m^6^A writers and m^6^A eraser found in early-diverging fungi and species from the Pucciniomycotina (up to five m^6^A writers in a species), not many m^6^A readers containing an YTH domain were identified, suggesting that there may be different types of RNA-binding proteins that recognize m^6^A modification. It would be interesting to study possibly divergent roles of m^6^A factors in these relatively understudied fungal taxa.

### Expression levels of m^6^A factors throughout the life cycle of *F. graminearum*

The levels of *MTA1* and *YTH1* expression were significantly elevated at 15 minutes after placing conidia on Bird medium (S1) and then decreased sharply to a basal level during polar growth and hyphal branching (S2–S3). These findings suggest that the potential m^6^A writer and m^6^A reader likely have a role during the initial stage of conidial germination. In line with this hypothesis, we observed slower conidial germination and less hyphal branching in the *MTA1*-OE5 strain. Although m^6^A demethylase activities of *ALKBH* genes have never been confirmed in fungi, the three putative m^6^A erasers, *ALKBH1*, *ALKBH4*, and *ALKBH5*, were induced during polar growth (S2) or hyphal branching (S3). These possible m^6^A erasers might be involved in maintaining low m^6^A levels during active vegetative growth. The significant increase in expression of actin and tubulin genes at S2– S3 reflected extensive hyphal growth and branching. Specifically, the expression level of *ALKBH4* reached its peak at S0 (fresh conidia), but experienced a sharp decline at S1 (the initial stage of conidial germination). However, it was gradually induced during polar growth and hyphal branching stages (S2-S3), exhibiting an opposite expression pattern compared to *MTA1* and *YTH1*. Although we did not observe any phenotypic changes in the knockout mutant lacking *ALKBH1*, it would be possible that other *ALKBH* genes may be involved in the regulation of conidial germination and hyphal growth by maintaining low m^6^A level in *F. graminearum*. Among the five putative m^6^A erasers, *ALKBH2* showed dramatic increase in expression during the ascospore formation. These expression dynamics of m^6^A factors highlight the complex regulation and involvement of m^6^A factors throughout different stages of the *F. graminearum* life cycle.

### Concluding remarks

The largest subphylum Pezizomycotina in fungi underscores their importance and their complex interactions with humans, encompassing a wide range of fungi that have significant influences on humans, both negative and positive. Examples include the opportunistic human pathogen *Aspergillus fumigatus* and the dermatophyte *Trichophyton rubrum*, as well as many plant pathogenic fungi causing serious diseases on our staple crops, such as *F. graminearum* and *M. oryzae*. The Pezizomycotina also includes several ecologically significant species, encompassing those that play crucial roles in wood and litter decay processes, as well as those that form symbiotic associations with other organisms, including lichens. In filamentous fungi, the m^6^A writer *MTA1* appears to have evolved to have important roles in adapting to diverse ecological niches, particularly in relation to filamentous growth. To identify target genes of *MTA1* that caused delayed conidial germination and slower hyphal growth in the *MTA1*-OE5 strain, we are under investigation for potential m^6^A sites in transcripts, using an Oxford Nanopore direct RNA sequencing technology.

## Materials and methods

### Phylogenetic analyses of m^6^A factors in fungi

We downloaded protein sequences from the UniProt database (https://www.uniprot.org/) that possessed pfam domains associated with m^6^A factors. These domains include PF05063 (MT-A70), PF04146 (YTH), PF13532 (ALKB), PF17098 (WTAP/MUM2), and PF15912 (Virilizer, N-terminal). However, the initial lists of potential m^6^A factors contained duplications and likely misannotated sequences. To address this issue, we conducted a manual inspection of the lists, excluding sequences that were either too short or too long, as well as sequences with small *E* -values, applying specific thresholds (e.g. *E*-value > 10-5). Whenever a sequence was removed from the lists, we also eliminated all entries from the corresponding species to ensure an accurate estimation of the number of m^6^A factors per species. The seqeunces for each pfam domain were aligned using MAFFT (v7.310) with the ‘auto’ setting (72). Poorly aligned regions of the resulting multiple sequence alignment were trimmed, using the Trimal program, with the parameter setting ‘– gappyout’ (73). To determine the best protein substitution model for each pfam domain, we used a perl scrip that can be found in the following website (https://github.com/stamatak/standard-RAxML/blob/master/usefulScripts/ProteinModelSelection.pl). We selected protein substitution models GAMMALG for m^6^A writers, GAMMAJTTF for m^6^A readers, GAMMALGF for ALKBHs and VIRN, and GAMMAJTT for WTAP. Maximum likelihood trees were constructed using the RAxML program (v8.2) (74). Outgroup was set to the homologue of m^6^A factors in *Arabidopsis thaliana* and nodal supports were evaluated by 1,000 bootstrap replications.

### Genetic transformation for gene deletion and overexpression

To generate gene deletion mutants in *F. graminearum* PH-1 strain, we employed a split marker strategy (75). This involved amplifying the left and right flanking regions of the target genes and combining them with a minimal gene cassette that carried the hygromycin phosphotransferase (*HPH*) gene under the control of the *trpC* promoter from *Aspergillus nidulans*. To achieve this, we conducted fusion PCR, following the previously described method (76), and the specific primers used for targeted gene deletion can be found in Supplementary Table S3. In brief, we amplified the left and right flanking regions of the target genes separately, using L5 and L3 primer pairs and R5 and R3 primer pairs, respectively. The L3 and R5 primers contained 27-nucleotide (nt)-long overhang sequences that were complementary to the 5’ and 3’ ends of the minimal *HPH* cassette (1,376-bp in length). The *HPH* cassette was obtained from the pCB1004 plasmid (77), amplified using HYG-F and HYG-R primers. Subsequently, we merged the PCR amplicons through overlap extension, assembling the left flanking region and the *HPH* cassette, or the right flanking region and the *HPH* cassette. Th split marker constructs were obtained by amplifying the fused amplicons using nested primer pairs (N5 and HY-R primers for the left-half construct and YG-F and N3 primers for the right-half construct). Finally, we introduced the two split marker constructs into protoplasts through polyethylene glycol-mediated transformation (78). Following transformation, transformants resistant to 200 mg/mL of hygromycin were examined for replacement of the target gene with the *HPH* cassette by diagnostic PCR checks, in which L5 and R3 primers that aneal to flanking seqeunces of the homologous recombination event were used to confirm homologous integration of the *HPH* cassette into the target loci.

For generation of *MTA1*-overexpressing strains, the coding seqeunce of *MTA1* was cloned to pDS23 plasmid digested with BglII and HindIII resctriction enzymes (79), using the In-Fusion HD Cloning kit (Takara Bio, Otsu, Japan). Five micrograms of the plasmid harboring *MTA1* were transformed into protoplasts of the WT strain. Following transformation, transformants resistant to 200 mg/mL of nourseothricin (Jena Bioscience, Jena, Germany) were examined for introduction of an additional copy of *MTA1* by diagnostic PCR check using primer pair, MTA1_OE_RT_fwd and MTA1_RT_rev. Primers used in this study are listed in Supplementary Table S3.

### Sexual development and hyphal growth measurement

Carrot agar plates (60 mm in diameter) (80) were inoculated by placing an agar block containing hyphae of *F. graminearum* at the center. The plates were incubated at room temperature under constant light. Six days after incubation, mycelia were removed by gently scraping the surface with a spatula, and then 0.9 ml of 2.5% Tween 60 (Sigma-Aldrich, St. Louis, MO, USA) was applied to the surface to assist the formation of perithecia. Sexual development in knockout mutants was observed by a stereomicroscopy for size and number formed. Squash mounts of young developing perithecia in water were examined using a compound microscope to check morphology and maturity of ascospores.

To obtain macroconidia of the WT and *MTA1*-overexpressing strains, small agar blocks containing each strain were inoculated and cultured in carboxymethylcellulose medium for 3 days at 20°C (81). Macroconidia were harvested by filtration through two layers of Miracloth. Concentrations of macroconidia were adjusted to 1 × 10^8^ spores per mL, and 40 μL of spore suspensions were inoculated into 100 mL of q-PDB. Cultures were shaken at 150 rpm at room temperature. Mycelia were collected by filtration through one layer of Miracloth and oven-dried at 55^◦^C for two days.

### m^6^A quantification

The m^6^A RNA methylation level was assessed using the EpiQuikTM m6A RNA methylation quantitative kit (Epigentek, Farmingdale, NY, USA). Briefly, 200 ng of total RNA was added and bound with antibody in the strip wells, followed by the process of washing, capture and detection of antibody. Finally, the signals were detected colorimetrically by reading the absorbance at 450 nm. m^6^A levels were eventually calculated based on a constructed standard curve.

### RT-PCR and quantitative RT-PCR analyses

Total RNA was extracted from frech conidia, hyphae and perithecial tissues ground in liquid nitrogen using TRIzol reagent (Thermo Fisher Scientific) according to the manufacturer’s instructions together with the following extraction steps: two phenol (pH 4.6)-chloroform-isoamyl alcohol (25:24:1) extraction steps followed by two chloroform extraction steps after the initial TRIzol-chloroform phase separation. RNA pellets were dissolved in 88 μL of nuclease-free water and subjected to genomic DNA digestion with DNase (Qiagen Inc.). RNA samples were then concentrated using RNA Clean & Concentrator (Zymo Research). For reverse transcription (RT)-PCR analysis of *MTA1*, 200 ng of RNAs were reverse transcribed and amplified using the OneStep RT-PCR kit (Qiagen Inc.). *MTA1* and *EF1α* (FGRRES_08811) were amplified for 34 cycles in the experiment for Figure 4C, or for 30 cycles in the experiment for Figure 5A, respectively. Annealing temperature was 62°C. The primer set for amplifying *EF1α* was designed to amplify flanking exons, including two introns in order to detect possible genomic DNA contamination. For semi-quantitative RT-PCR assays, first-strand cDNA synthesis was prepared from 200 ng of total RNA using the iScript cDNA Synthesis Kit (Bio-Rad), according to the manufacturer’s instructions. Real-time RT-PCR analyses were performed using the CFX96 Touch

Real-Time PCR Detection System (Bio-Rad). RT-PCR mixtures were composed of 1.5 μl of each primer (10 μM), 5 μl of SYBR Green Supermix (Bio-Rad), and 2 μl of cDNA (100 ng/mL). The PCR conditions consisted of an initial denaturing step at 95°C for 3 min, a denaturation step at 95°C for 10 s, both annealing and extension steps at 65°C for 30 s for 40 cycles, and 65 to 95°C with a 0.5°C increment and each temperature for 5 s to obtain the melting curve. The quantification of the relative expression levels was performed with the comparative cycle threshold method normalization (82), in that the expression level of *MTA1* was normalized against the reference gene *EF1α*. The averages of the three biological replicates and standard deviations of the relative expression values were presented (Supplementary Fig. S4). Primers used in the expression analysis are listed in Supplementary Table S3.

### RNA-seq and differential expression analyses

Total RNA samples were extracted from fresh conidia of the WT, *MTA1*-KD1, and *MTA1*-OE5 strains that have been harvested from carboxymethylcellulose medium 4 days after incubation at 20°C (81). Three separate experiments were performed to produce conidia, and the conidia samples from each experiment were used as replicates for RNA-seq analysis. Two micrograms of total RNA were sent to Macrogen Inc. (Seoul, Korea) for cDNA library construction, using the TruSeq Stranded Total RNA Library Prep Gold Kit (Illumina, San Diego, CA, USA), and for sequencing on the HiSeq4000 platform (Illumina). Raw reads (paired-end, 100 bp) were further processed and filtered, using the TrimGalore (v0.6.6) (https://www.bioinformatics.babraham.ac.uk/projects/trim_galore/). Filtered reads were mapped to the genome sequence of *F. graminearum* (NCBI accession: GCA_000240135.3), using the HISAT2 program (v2.1.0). The gene annotation file used in this study was Ensembl annotation v.32 (83). Mapped reads on genomic features, such as exon and intron, were calculated, using the htseq-count program. Gene expression levels in reads per kilobase per million mapped reads (RPKM) values were computed and normalized by effective library size estimated by trimmed mean of M values, using the edgeR R package (v3.26.8). For differential expression analysis, only genes with CPM values greater than 1 in at least 3 samples were kept for further analyses (9,792 out of 16,001 gene loci). Then, differentially expressed (DE) genes showing greater than 4-fold difference at an FDR of 5% were identified between of the WT, *MTA1*-KD1, and *MTA1*-OE5 strains, using the limma R package (v3.28.21).

### DATA accessions

The RNA-seq data generated in the present work have been deposited in NCBI’s Sequence Read Archive and are accessible through SRA accessions from SRX21157089 to SRX21157097, which belong to the BioProject (accession, PRJNA998561). RNA-seq data for investigating the expression level changes in m^6^A factors throughout the life cycle of *F. graminearum* can be found in NCBI’s Gene Expression Omnibus GSE109088 for the conidial germination stages (S1–S3) and GSE109094 for the sexual development (S4–S9).

### Availability of data and material

All data generated or analyzed during this study are included in this published article and its supplementary information files.

## Competing interests

The authors declare that they have no competing interests.

## Acknowledgements

This publication is based upon work supported by the Basic Science Research Program through the National Research Foundation of Korea, funded by the Ministry of Education (2019R1I1A1A01057502). The funders had no role in study design, data collection and analysis, decision to publish, or preparation of the manuscript.

## Authors’ contributions

HK performed functional analyses of m^6^A factors. JH and HK conducted m^6^A quantification. WK conducted phylogenetic analyses of m^6^A factors. HK and WK analyzed and interpreted the data. HK and WK wrote the manuscript. All authors read and approved the final manuscript.

## Supplemental Material

Supplemental material is available online only

Figure S1. Maximum likelihood Trees of potential m^6^A factors in fungi

Figure S2. Normal ascospore production in m^6^A factor mutants.

Figure S3. Genotyping for *MTA1*-deletion mutants.

Figure S4. *MTA1* overexpression and knockdown.

Figure S5. Dynamic expression patterns of potential m^6^A eraser genes.

Figure S6. Reduced hyphal branching in the *MTA*1-OE5 strain.

Figure S7. MA plots for differential expression analyses.

Table S1. Metadata for UniProt protein sequences related to MTA1, IME4, KAR4, WTAP, VIRN, YTH1 and ALKBHs.

Table S2. Numbers of putative m^6^A erasers found in 27 selected fungi.

Table S3. Primers used in this study.

